# Genome Assembly of the Popular Korean Soybean Cultivar Hwangkeum

**DOI:** 10.1101/2021.04.19.440529

**Authors:** Myung-Shin Kim, Taeyoung Lee, Jeonghun Baek, Ji Hong Kim, Changhoon Kim, Soon-Chun Jeong

## Abstract

Massive resequencing efforts have been undertaken to catalog allelic variants in major crop species including soybean, but the scope of the information for genetic variation often depends on short sequence reads mapped to the extant reference genome. Additional *de novo* assembled genome sequences provide a unique opportunity to explore a dispensable genome fraction in the pan-genome of a species. Here, we report the *de novo* assembly and annotation of Hwangkeum, a popular soybean cultivar in Korea. The assembly was constructed using PromethION nanopore sequencing data and two genetic maps, and was then error-corrected using Illumina short-reads and PacBio SMRT reads. The 933.12 Mb assembly was annotated 79,870 transcripts for 58,550 genes using RNA-Seq data and the public soybean annotation set. Comparison of the Hwangkeum assembly with the Williams 82 soybean reference genome sequence revealed 1.8 million single-nucleotide polymorphisms, 0.5 million indels, and 25 thousand putative structural variants. However, there was no natural megabase-scale chromosomal rearrangement. Incidentally, by adding two novel groups, we found that soybean contains four clearly separated groups of centromeric satellite repeats. Analyses of satellite repeats and gene content suggested that the Hwangkeum assembly is a high-quality assembly. This was further supported by comparison of the marker arrangement of anthocyanin biosynthesis genes and of gene arrangement at the *Rsv*3 locus. Therefore, the results indicate that the *de novo* assembly of Hwangkeum is a valuable additional reference genome resource for characterizing traits for the improvement of this important crop species.

## Introduction

Hwangkeum is an important soybean [*Glycine max* (L.) Merr.] cultivar with distinctive organoleptic and agronomical features. Ever since its cultivar release in 1979 (Park *et al*. 1981), it has been widely grown and widely used as a breeding parent in Korea. According to the 2008 national survey report (Yu *et al*. 2008), it was used as a parent or grandparent in 19 of the 105 newly bred soybean cultivars released in Korea up to 2007. Hwangkeum has a determinate growth habit and non-shattering pods, and is adapted to the middle Korean peninsula (Maturity Group V). Seeds are large (25 g per 100 seeds), round-shaped, and clear golden with yellow seed-coats and buff hila (Yang *et al*. 2010). Hwangkeum was found to be resistant to all soybean mosaic virus (SMV) strain groups identified in the USA (Chen *et al*. 2002), and the resistance was found to be conferred by multiple genes (Jeong and Jeong 2014). The genes controlling anthocyanin biosynthesis are highly polymorphic between Hwangkeum and IT182932, a wild soybean accession (Yang *et al*. 2010). Low isoflavone content in Hwangkeum led to the identification of novel loci that regulate the content of isoflavone (Yang *et al*. 2011).

The first genome sequence of soybean, one of the major seed crop species worldwide, was that of Williams 82, which was published in 2010 (Schmutz *et al*. 2010). The Williams 82 soybean reference genome sequences were generated using a whole-genome shotgun approach with Sanger-sequencing, and then assembled with physical and high-density genetic maps. Subsequently, additional genome assemblies that were supposed to represent soybean growing areas have been generated with high-throughput sequencing platforms: Japanese cultivar Enrei (Shimomura *et al*. 2015), Chinese cultivar Zhonghuang 13 (Shen *et al*. 2018), and southern US cultivar Lee (Valliyodan *et al*. 2019), while classifying Williams 82 as a northern US cultivar. Additionally, the genome sequences of two wild soybean accessions W05 (Xie *et al*. 2019) and PI 483463 (Valliyodan *et al*. 2019), and of a perennial relative of soybean, *Glycine latifolia* (Liu *et al*. 2018), have already been published. These efforts have recently culminated in the construction of a high-quality pan-genome from 26 diverse soybean accessions sequenced individually using single molecule real-time (SMRT) sequencing, together with the existing Williams 82, Zhonghuang 13, and W05 genomes (Liu *et al*. 2020).

Degrees of structural variation of these genome sequences from that of Williams 82 are highly variable. For example, comparison between those of Williams 82 and Zhonghuang 13 revealed many putative mega-scale structural variants, while none were observed between those of Williams 82 and Lee. Here, we report our investigation of the Hwangkeum genome using PromethION nanopore sequencing data and two genetic maps. We show that most of the mega-scale structural variants between Hwangkeum and Williams 82 assemblies might be assembly errors. Besides those mega-scale variations, most of the small and structural variants between the two genome assemblies might be natural. The observed differences were validated by examination of known variation regions, including anthocyanin biosynthesis genes and disease resistance genes.

## Materials and Methods

### Plant materials and sequencing

Seeds of Hwangkeum whose breeding line was known as Suwon 97 (Chen *et al*. 2002; Jeong and Jeong 2014) were planted in the greenhouse at the Korea Research Institute of Bioscience and Biotechnology. After three weeks’ growth, a bulk of young trifoliolate leaf tissues was collected for genomic DNA extraction. Note that the seeds of Hwangkeum used in this study came from the line of Hwangkeum that had been subject to single plant selection at least twice during our recent 180K SoyaSNP array and genome resequencing studies (Lee *et al*. 2015; Kim *et al*. 2021). Genomic DNAs for the generation of Illumina short-read (Illumina, San Diego, CA, USA) and PacBio SMRT long-read sequences (Pacific Biosciences, Menlo Park, CA, USA) were extracted using the CTAB method, as described by Saghai-Maroof *et al*. (1984). Paired-end and mate-pair libraries for Illumina short-read sequencing were prepared, and then sequenced mainly using a HiSeq 2500 System. A library for PacBio SMRT sequencing was prepared using SMRTbell Express Templates with Sequel SMRT Cell 1M v2, Sequel Binding Kit 2.1, and was then sequenced with a PacBio Sequel system. Genomic DNA for the single-molecule sequencer PromethION (Oxford Nanopore Technologies Ltd., Oxford, UK) sequencing was extracted using Nanobind Plant Nuclei Big DNA Kit -Alpha Version (#NB-900-801-01) (Circulomics Inc., Baltimore, MD), as described by Workman *et al*. (2018), and was further purified using 26G Needle shearing and Bluepippin size selection (High Pass Plus, (20 - 150) kb). The purified DNA was then prepared for sequencing following the protocol in the genomic sequencing kit SQK-LSK109 (Oxford Nanopore Technologies Ltd.).

For the extraction of total RNAs, plants were further grown to a pod-bearing stage, and the bulks tissues were separately collected. Total RNAs were extracted from the six different tissues using RNeasy Plant Mini Kit, following the manufacturer’s instructions (QIAGEN, Venlo, Netherlands). Two separately combined RNA extracts were used for RNA sequencing (RNA-seq). Equal amounts of the RNAs extracted from immature seeds, young shoot, and young stems were combined into one sample, and the RNAs from flowers, leaves, and roots were combined to form another sample. Libraries for each of the RNA samples were prepared using TruSeq RNA Sample Prep Kit v2 (Illumina), and then 101 bp paired-end short reads were generated on an Illumina platform.

### Genome assembly

*PacBio SMRT data* Assembly of SMRT subreads was performed with FALCON-Unzip to produce primary contigs (Chin *et al*. 2016). The primary contigs were polished with mapped PacBio subreads with Quiver implementation in variantCaller tool (SMRT Link 6.0.0.47841; https://www.pacb.com/support/software-downloads/) with three iterations, followed by. Pilon (v1.22) (Walker *et al*. 2014) with Illumina data. Mate-pair reads were used to construct scaffolds with the SSPACE program (v2.3.1) (Boetzer *et al*. 2011), with sequence gaps filled with PBJelly (v15.8.24) (English *et al*. 2014). The scaffolding and gap-filling were then repeated with paired-end reads. Finally, ALLMAPS (Tang *et al*. 2015) was used to construct the 20 pseudo-chromosomes by anchoring the assembled contigs/scaffolds to two genetic maps (WH and HI maps) that had been constructed using Hwangkeum as a parental line (Lee *et al*. 2020). In our previous study, we constructed four high-density genetic maps from Williams 82K (*G. max*) by Hwangkeum (*G. max*) (referred to as WH), Hwangkem by IT182932 (*Glycine soja*) (HI), Williams 82K by IT182932 (WI), and IT182932 by IT182819 (*G. soja*) (II) populations. To remove missing markers in the assemblies, probe or primer sequences of markers were searched against the assembly using BLAST+ (Camacho *et al*. 2009), and the marker sequences hit by > 95% identity and > 88% coverage were input into the ALLMAPS program, with equal weight assigned to the two genetic maps. *Nanopore PromethION data* All PromethION reads were assembled into contigs with Shasta v.0.1.0 (Shafin *et al*. 2020) to obtain raw genome assembly results. Then, ALLMAPS (Tang *et al*. 2015) was used to construct the 20 pseudo-chromosomes, as described above. The resulting assemblies were polished with Pilon (v1.22) (Walker *et al*. 2014) with three iterations with mapping of Illumina short reads, and with Arrow implemented SMRT Link 8.0.0.80529 with three iterations with mapping of SMRT reads. To assess the completeness of the final genome, Benchmarking Universal Single-Copy Orthologs (BUSCO) (Simão *et al*. 2015) was employed using eukaryota odb10 (creation date: November 20, 2019, number of species: 70, number of BUSCOs: 255) and embryophyta odb10 (creation date: November 20, 2019, number of species: 50, number of BUSCOs: 1614) core conserved genes as databases.

### Comparative genomics between Williams 82 and Hwangkeum

We identified SNPs and indels (< 50 bp) using paftools.js from the minimap2 distribution (Li 2018). Briefly, we mapped the Hwangkeum assembly as a query against the Williams 82 (Wm82.a2.v1) assembly as a reference using minimap2, and called variants through the paftools.js module in minimap2 with the following flags (minimap2 -c --cs ref.fasta query.fasta | sort -k6,6 -k8,8n | paftools.js call -L15000).

We identified and classified the structural variants using the Structural Variants from MUMmer (SVMU) pipeline (Chakraborty *et al*. 2018; Marçais *et al*. 2018). Insertion (INS) or deletion (DEL) was classified on the basis of whether the Hwangkeum assembly had longer or shorter sequence, respectively, with respect to the reference genome Williams 82 sequence. Translocation and inversion events (both refer to structure variation ≥ 1.0 Kbp) were detected by manual check depending on their location and orientation to their neighboring blocks, based on the non-allelic homology blocks from the above alignment, using MUMmer4 (v. 4.0.0beta2) (Marçais *et al*. 2018).

Visual evaluations for structural comparisons between assemblies were made using dot plots generated by the MUMMERPLOT utility from MUMMER v.4.0 (Marçais *et al*. 2018). Correspondences of orthologous genes between Hwangkeum and Williams 82 were determined using OrthoMCL (v2.0.9) with default options (Li *et al*. 2003). We used the MCscan (Python version) (Tang *et al*. 2008) to compare gene arrangement at the *Rsv*3 locus between the Hwangkeum and Williams 82 assemblies.

### Analysis of telomeric and centromeric repeats

As a measure of pseudomolecule completeness near the chromosome ends, we checked for characteristic telomeric repeat motifs AAACCCT and AGGGTTT within 1,500 bases of the leading and trailing ends of the pseudomolecule ends (Valliyodan *et al*. 2019). Additionally, we searched for any novel repeat elements in the terminal sequences with Tandem Repeats Finder (Benson 1999).

We searched for two centromere-specific satellite repeats (CentGm-1 and CentGm-2), which have been predicted using sequencing data (Vahedian *et al*. 1995; Swaminathan *et al*. 2007; Gill *et al*. 2009; Tek *et al*. 2010), and then confirmed experimentally (Gill *et al*. 2009; Findley *et al*. 2010), in order to identify the assembled centromeric regions in the Hwangkeum and Williams 82 assemblies. Representative consensus sequences of CentGm-1 and -2 were proposed from the analysis of three soybean assemblies by Valliyodan *et al*. (2019). These representative satellite repeat consensus sequences were aligned with the Williams 82 and Hwangkeum assemblies with an -evalue 1e-5 -task blastn-short -penalty -1 option in BLASTN to estimate the location and length of the centromeres on the pseudomolecules. All the repeat sequences hit by each of the CentGm-1 and -2 sequences had > 67% sequence identity with their query sequences. We then further filtered these candidate repeats with < 80% alignment coverage. Note that < 80% alignment coverage and < 60% sequence identity were a cut-off criteria used in a previous phylogenetic analysis of a whole-genome shotgun database (Gill *et al*. 2009). A majority of repeat sequences hit by each of the CentGm-1 and -2 sequences appeared to overlap each other, likely due to the 81.5% sequence identity between the CentGm-1 and CentGm-2, and thus the two extracted sequence sets for each of the Hwangkeum and Williams 82 assemblies were combined into a set of repeat sequences by removing one of the overlapped sequences. Lengths of the satellite tandem repeats in pseudomolecules and unanchored contigs were determined with the Tandem Repeat Finder (Benson 1999).

The combined repeats from the Hwangkeum assembly were further filtered for efficient phylogenetic analysis. First, 4,599 repeats with length < 89 bp and 38 with > 94 bp were excluded. The cd-hit-est software was then used to cluster similar repeat sequences into clusters using the parameters “-c 0.90 -n 10” within a set of 20,386 satellite repeats (Fu *et al*. 2012). Multiple sequence alignment of the resultant non-redundant 4,469 satellite repeats was performed with ClustalW (Larkin *et al*. 2007), and then phylogenetic analysis of the aligned sequences was performed with MEGA7 software using the neighbor-joining method (Kumar *et al*. 2016). In this phylogenetic analysis, four CentGm-1 (referred to as CentCm-1_AF, CentCm-1_E, CentGm-1_Gill, and CentCm-1_J2), three CentGm-2 (CentCm-2_G, CentCm-2_Gill, and CentCm-2_M) representative sequences used for karyotyping soybean by Findley *et al*. 2010, and two (CentGm-1_V and CentGm-2_V) consensus sequences proposed by Valliyodan *et al*. 2019 were included as reference sequences to infer the already established CentGm-1 and CentGm-2 repeat groups.

### Genome annotation

Repetitive sequences were identified with RepeatMasker (v. 4.1.1; http://repeatmasker.org) with -s -pa 15 -no_is -xsmall -gff -lib options using a soybean repeat library from SoyTEdb (Du *et al*. 2010). We annotated gene models using the Seqping pipeline (Chan *et al*. 2017) with slight modifications. Seqping uses transcriptome data and three self-training Hidden Markov Model (HMM) models, and the resultant predictions are then combined using MAKER2 (Holt and Yandell 2011). We added protein models at the MAKER2 step. The predicted genes were filtered out using e-AED value with threshold of 0.4. For the transcript data to train the prediction models, we used RNA-seq data generated from the Hwangkeum tissues described above. The RNA-seq data were processed with genome-guide assembly, and gene structures were then predicted by the EMBOSS getorf program with the default parameters. All the resultant gene model sets were integrated into single RNA-seq-based gene model sets. *Glycine max* protein set downloaded from NCBI database was used as a reference protein file for the validation and annotation of the gene predictions. We used tRNAscan-SE software (version 2.0) with default parameters for tRNA annotation (Chan and Lowe 2019) and Barrnap 0.9 (https://github.com/tseemann/barrnap) for rRNA annotation. Protein function annotations were added by searching for homologous proteins in the UniProt SwissProt database (Bateman *et al*. 2017) using BLASTP and eggNOG v4.5 database (Huerta-Cepas *et al*. 2016) using psi-blast with E-value < 1e-5, num_alignments 5, and num_descriptions 5, and protein domains using InterProScan 5.34-73.0 (Finn *et al*. 2017). The functional annotation results were read using Annie (http://genomeannotation.github.io/annie/), and then genome annotation summary statistics were generated using the software GAG (Geib *et al*. 2018).

Nucleotide-binding and leucine-rich-repeat (NLR) genes, which are members of the largest resistance gene family in plants, were predicted using TGFam-Finder (v. 1.03) (Kim *et al*. 2020). TGFam-Finder is a domain search-based gene annotation tool. We used the NB-ARC domain (PfamID = PF00931) (van der Biezen and Jones 1998), which was used in the TGFam-Finder program, as TARGET_DOMAIN_ID for searching NLR genes. Transcriptome mapping was performed using the RNA-seq data generated from the Hwangkeum tissues described above. We searched for only primary transcripts from the Hwangkeum genome sequence.

## Data availability

All whole genome sequencing data are available at NCBI (Bioproject PRJNA628825) except a set of paired-end short reads downloaded from NCBI with accession number: SRX6472178. The genome assembly and annotation data of Hwangkeum v.1.0 is deposited at GenBank under the accession JAGRRG000000000. Supplemental material (Figures S1-S5 and Table S1-S10) and six supplemental data Files are available at Figshare. The supplemental data Files are two Tandem Repeat Finder results (File S1 and File S4), SNPs and indels (File S2), structural variants (File S3), a list of annotated transcripts (File S5), and a list of NLR genes (File S6).

## Results and Discussion

### Genome assembly of the Hwangkeum

The genome of *Glycine max* cv. Hwangkeum was sequenced at 78× coverage (78,861,723,603 bases) using PacBio SMRT technology, and at 89× coverage (89,519,105,740 bases) using Nanopore PromethION technology. Both the sequencing data were separately assembled with error corrections up to pseudomolecules. The diploid FALCON-Unzip assembler produced an initial SMRT-based contig assembly with 1,436 primary contigs, N50 of 1.71 Mb, and a total length of 963.13 Mb (Table S1). After error corrections and scaffolding using Illumina mate-pair and paired-end reads, the final primary assembly was scaffolded into 730 scaffolds covering 966.25 Mb with an N50 of 2.54 Mb and with a maximum length of 11.72 Mb (Table S2). We initially evaluated two recently published assemblers, Shasta and wtdbg2 (Ruan and Li 2020; Shafin *et al*. 2020), on our PromethION read data (Table S1). Total lengths of both the assemblies from the PromethION data were approximately 30 Mb shorter than that from the SMRT data. The Shasta assembly showed approximately 8 times fewer number of contigs (847) and 10 times longer N50 length (6.95 Mb) relative to those of the wtdbg2 assembly. Thus, the results showed that, despite much higher levels of differences, the tendency was somewhat consistent with that from the human genome assembly study (Shafin *et al*. 2020), suggesting that Shasta might be more appropriate than wtdbg2 for the assembly of our Hwangkeum PromethION sequencing data.

To further evaluate which of the FALCON-Unzip SMRT and Shasta PromethION assemblies was superior, we then generated chromosome-scale pseudomolecules by ordering and orienting the assembled contigs/scaffolds via anchoring to two genetic maps that had been constructed using Hwangkeum as a parental line (Lee *et al*. 2020). Our comparison between four genetic maps, including the two Hwangkeum genetic maps, showed excellent collinearity with no marker order difference, although there appeared to be putative megabase-scale inversions based on the lack of cross-overs. Thus, we hypothesized that the assembly that showed the lesser number of discrepant markers between sequence assembly and genetic maps was likely superior to the other. The final assembly of Hwangkeum on the SMRT data consisted of 944.02 Mb of 20 chromosome-level pseudomolecules containing 640 scaffolds and 22.32 Mb of 90 unplaced scaffolds, while that on the PromethION data consisted of 907.90 Mb of 20 chromosome-level pseudomolecules containing 399 scaffolds and 19.74 Mb of 448 unplaced contigs. Thus, approximately 30 Mb longer sequences of SMRT scaffolds relative to that of the PromethION scaffolds were anchored to 20 chromosome-scale pseudomolecules. For the SMRT pseudomolecules, 553.39 Mb of 201 scaffolds were oriented with more than four markers, while 634.65 Mb of 90 contigs for the PromethION pseudomolecules were well oriented (Table S3), suggesting that the approximately 80 Mb sequence was better oriented in the PromethION assembly than in the SMRT assembly. We then examined the number of translocation errors, which represent breaks in collinearity between sequence and genetic maps markers due to the mixing of non-homologous chromosomes as well as of the assembled pseudomolecules, in order to assess the integrity of scaffolds or contigs. From the SMRT pseudomolecule assembly, we observed 45 single-marker inter-chromosomal translocation errors, 121 multiple marker chimeric scaffolds with mappings to multiple linkage groups, and one apparent intra-chromosomal translocation on chromosome 13. In stark contrast, we observed only one chimeric scaffold on chromosome 18 from the PromethION pseudomolecule assembly. Three markers at the top of chromosome 18 appeared to best match with three different regions on chromosome 11. The results indicated that the PromethION-based assembly contained a much lower number of errors than the SMRT-based assembly in this study. Thus, we decided to use the PromethION-based assembly as a representative assembly of Hwangkeum genome in this study.

The initial PromethION-based assembly was then error-corrected using Pilon with the Illumina short reads and Arrow with the SMRT reads, which was a similar strategy to those used in the other plant genome assemblies (Xie *et al*. 2019; Jiao and Schneeberger 2020). When we mapped marker sequences from the WH and HI maps to the error-corrected assembly, we observed that the three markers at the top of chromosome 18 that best matched with the three different regions on chromosome 11 in the initial ALLMAPS assembly now best matched with the top region of chromosome 18. The final error-corrected Nanopore PromethION assembly had a total length of 933.12 Mb, and consisted of 913.20 Mb of 20 chromosome-level pseudomolecules containing 378 contigs and 19.92 Mb of 448 unplaced contigs (Table 1).

**Table 1.** Summary statistics of the Hwangkeum genome assembly.

| Assembly feature | Number | Size |
| --- | --- | --- |
| Total assembly length |  | 933,123,489 bp |
| Pseudomolecules | 20 | 913,200,796 bp |
| Unanchored contigs | 448 | 19,922,693 bp |
| Repetitive content |  | 468,186,948 bp (50.17%) |
| Centromeric satellite repeats | 25,030 | 2,249,110 bp (0.24%) |
| Number of transcripts | 79,870 |  |
| Number of genes | 58,550 |  |

### Evaluation of the assembly genome quality

Analyses with two BUSCO databases, eukaryota odb10 and embryophyta odb10, indicated that the genome content was effectively captured in the Nanopore PromethION assembly (Table S4): BUSCO analysis against eukaryota odb10 and embryophyta odb10 demonstrated 2/255 (0.7%) and 15/1,614 (0.9%) of BUSCO genes missing from the assembly, respectively. We found telomeric repeat motifs AAACCCT and AGGGTTT on only 9 of the 40 pseudomolecule ends in Hwangkeum relative to 23 in the Williams 82 reference sequence. The results indicated that although our PromethION sequencing is not nearly as efficient as Sanger shot-gun sequencing, it caught the ends of chromosomes.

We also evaluated distribution patterns of centromeric satellite repeats across chromosomes in the Hwangkeum assembly. Two groups of centromere-specific satellite repeat sequences (CentGm-1 and CentGm-2 with 92-bp and 91-bp monomers, respectively) have been reported using sequencing data (Vahedian *et al*. 1995; Swaminathan *et al*. 2007; Gill *et al*. 2009; Tek *et al*. 2010), and then confirmed by immunoprecipitation (Tek *et al*. 2010) and fluorescent *in situ* hybridization (Gill *et al*. 2009; Findley *et al*. 2010). Representative consensus sequences of CentGm-1 and -2 were recently proposed from the analysis of three soybean assemblies (Valliyodan *et al*. 2019), and thus we used these two sequences to identify the assembled centromeric regions in the Hwangkeum and Williams 82 assemblies. After filtration with a cutoff criterion of < 80% alignment coverage, we obtained 24,066 CentGm-1 and 22,046 CentGm-2 repeat sequences from the Hwangkeum assembly and 96,563 CentGm-1 and 92,749 CentGm-2 repeat sequences from the Williams 82 assembly. Thus, our cutoff threshold was less stringent than that used by Valliyodan *et al*. (2019) because they extracted only 11,829 CentGm repeats from the Williams 82 assembly. As expected from the 81.5% sequence identity between CentGm-1 and CentGm-2, a total of 21,612 repeat sequences were hit by both the query repeat sequences and thus their locations overlapped each other. Thus, the two extracted sequence sets from the Hwangkeum assembly were combined into a set of 25,030 repeat sequences (∼ 2.3 Mbp) (Table 1). Of the 25,030, the positions of 23,494 (93.8%) appeared to be head-to-tail tandem repeats, a feature typical of centromeric satellite repeats (Jiang et al. 2003). When their number, size, and locations were verified using Tandem Repeat Finder (Benson 1999), 24,859 (99.3%) of them appeared to be direct head-to-tail tandem repeats (Table S6 and File S1). The 91-bp CentGm-2 repeats were nearly absent (< 20 copies) on chromosome 18 and the 92-bp CentGm-1 repeats were absent on chromosomes 1 and 7 and nearly absent (< 20) on chromosomes 6, 9, 10 and 11. Thus, our results are somewhat consistent with a previous observation (Valliyodan *et al*. 2019) that copy numbers of identified tandem repeat units were highly variable between chromosomes, although this study showed wider distribution of the 91-bp CentGm-2 repeats across chromosomes unlike the previous observation. In the case of the Williams 82 assembly, we obtained a final combined set of 100,654 repeat sequences (∼9.2 Mb). Of the 100,654 repeats, 93,456 (92.8%) appeared to be head-to-tail tandem repeats. About 40.8% of the repeat sequences in the Hwangkeum assembly and ∼ 51.3% of them in the Williams 82 reference assembly were located in unanchored scaffolds, indicating that almost half of the highly repeated centromeric repeats were not incorporated into pseudomolecules. Our observation that the total numbers of centromeric repeats were approximately four times higher in the Williams 82 reference assembly than in the Hwangkeum assembly suggests that the assembly collapse of centromeric repeats is likely a main cause of the difference of total lengths of assemblies between Williams 82 and Hwangkeum (Tørresen *et al*. 2019).

### Genome structure comparison with other publicly available soybean genomes

Our recent genetic map study showed multiple mega-scale discordant regions between the Williams 82 reference genome and our genetic maps (Lee *et al*. 2020). However, comparison between the Williams 82 and Lee genome sequences resulted in no mega-scale structural variant (Valliyodan *et al*. 2019). In contrast, comparison between the Williams 82 and Zhonghuang 13 genome sequences identified many large (> 100 kb) structural variants (SV), including four mega-scale SVs (Shen *et al*. 2018, 2019). However, detailed investigation of whether the mega-scale SVs are real or miss-assemblies in either the assembly were not reported; neither did their subsequent pan-genome study address these mega-scale SVs (Liu *et al*. 2020). Thus, rather than comparing our Hwangkeum genome sequence and all other soybean *de novo* assemblies available, we decided in this study to focus on comparison between the current Hwangkeum and the Williams 82 reference genome.

Direct comparison between corresponding chromosome sequences of the Hwangkeum and Williams 82 assemblies identified 1,788,320 SNPs and 517,907 indels (< 50 bp) (Table S6 and File S2). Interestingly, the number of SNPs is similar to the combined number (1,678,164) of heterozygous (4,919), missing (713,953), and homozygous non-reference (959,292) SNPs for Hwangkeum in the 30,753,511 SNP set without the minor allele frequency filter detected in the 781 soybean haplotype map set (Kim *et al*. 2021). The number of indels is also similar to the combined number (470,389) of heterozygous (28,639), missing (303,784), and homozygous non-reference (137,966) indels for Hwangkeum in the 5,717,052 indel set without the minor allele frequency filtration detected from the same set. Thus, these observations suggested that the missing SNPs and indels might be real variants that were not easily detectable with short reads. Several chromosomal regions showed no difference between the Hwangkeum and Williams 82 assemblies. For example, the 85-cM gap in the middle of chromosome 4 for the WH population detected in our previous genetic mapping study (Lee *et al*. 2020) contained 15 no-variation regions of > 200 kb with the largest one of 1.72 Mb. These appear to be identity-by-descent regions inherited from a common ancestor during soybean breeding history.

In addition to the difference in the number and locations of centromeric repeats between the Hwangkeum and Williams 82 assemblies, most of the chromosomes in the Hwangkeum assembly were shorter in size, with a median decrease of 1.76 Mb, relative to corresponding chromosomes in the Williams 82 assembly (Figure 1A). Notable outliers were two of the greatest decreases that occurred in chromosomes 4 and 15, and increases observed in chromosomes 11 and 13. Aligning the Hwangkeum assembly to the Williams 82 assembly, we found additional notable megabase-scale rearrangements in these exceptionally decreased or increased chromosomes as well as in the other chromosomes (Figure 2B and Figure S1). All those mega-scale rearrangements located at the presumed pericentromeric regions where genetic markers are not resolved well due to low recombination rate. Interestingly, those exceptionally decreased or increased chromosomes could be explained by the insertions of unanchored scaffolds present in the Williams 82 assembly (chromosome 11) or by the corrected positioning of misjoints predicted by our genetic mapping study (chromosomes 4, 13, and 15), as described below.

**Figure 1.**
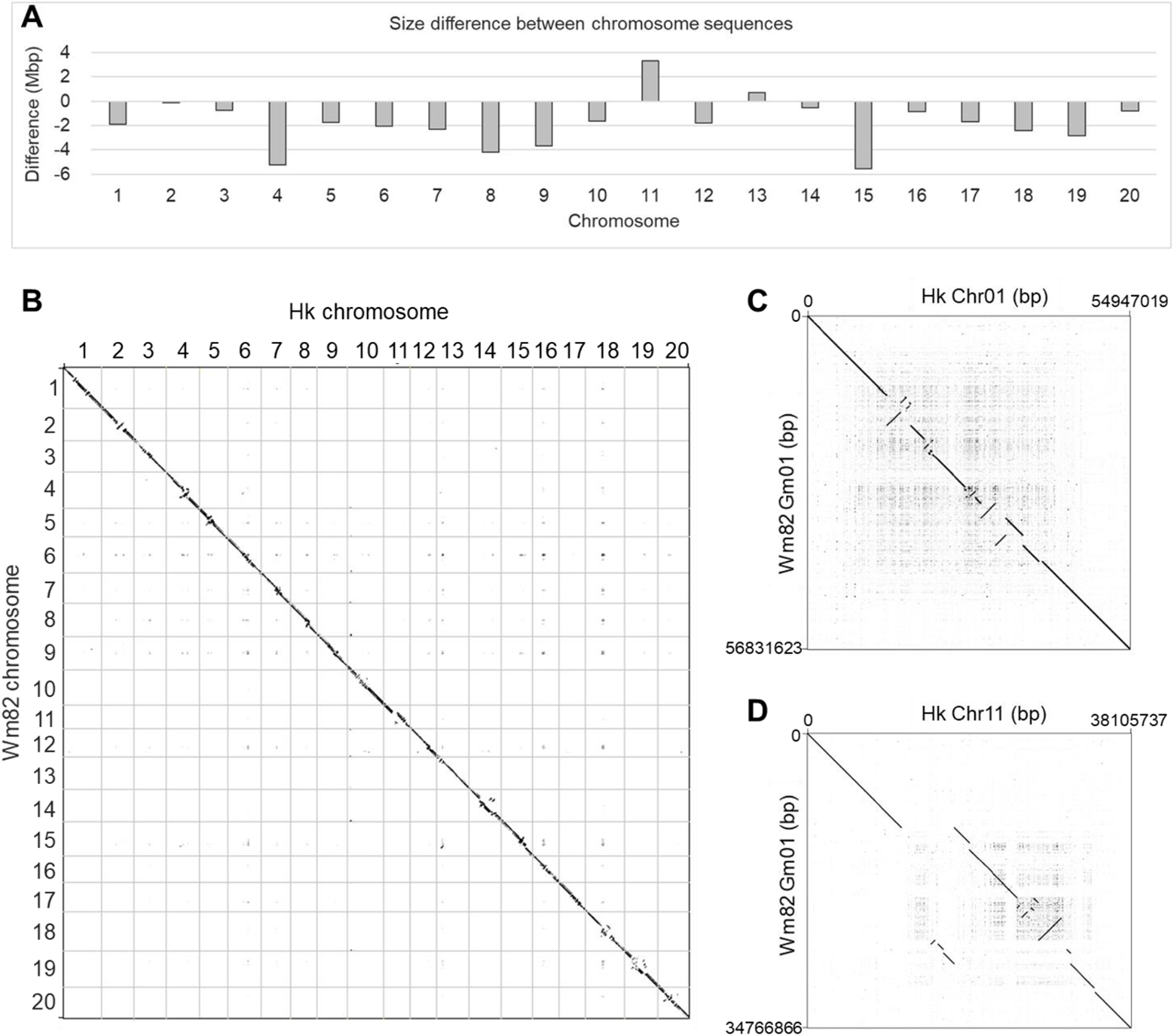
Comparison between the Hwangkeum and Williams 82 assemblies. A. Bar chart that shows size difference values between corresponding chromosomes of the Hwangkeum and Williams 82 assemblies. The values were obtained by subtracting length of each chromosome in the Williams 82 assembly from that of corresponding chromosome in the Hwangkeum assembly. B. Dot plots showing alignments of 20 chromosome sequences between the Hwangkeum (Hk) assembly and Williams 82 (Wm82) reference genome assembly and showing alignments of individual chromosomes 1 and 11 between the Hk and Wm82 assemblies.

**Figure 2.**
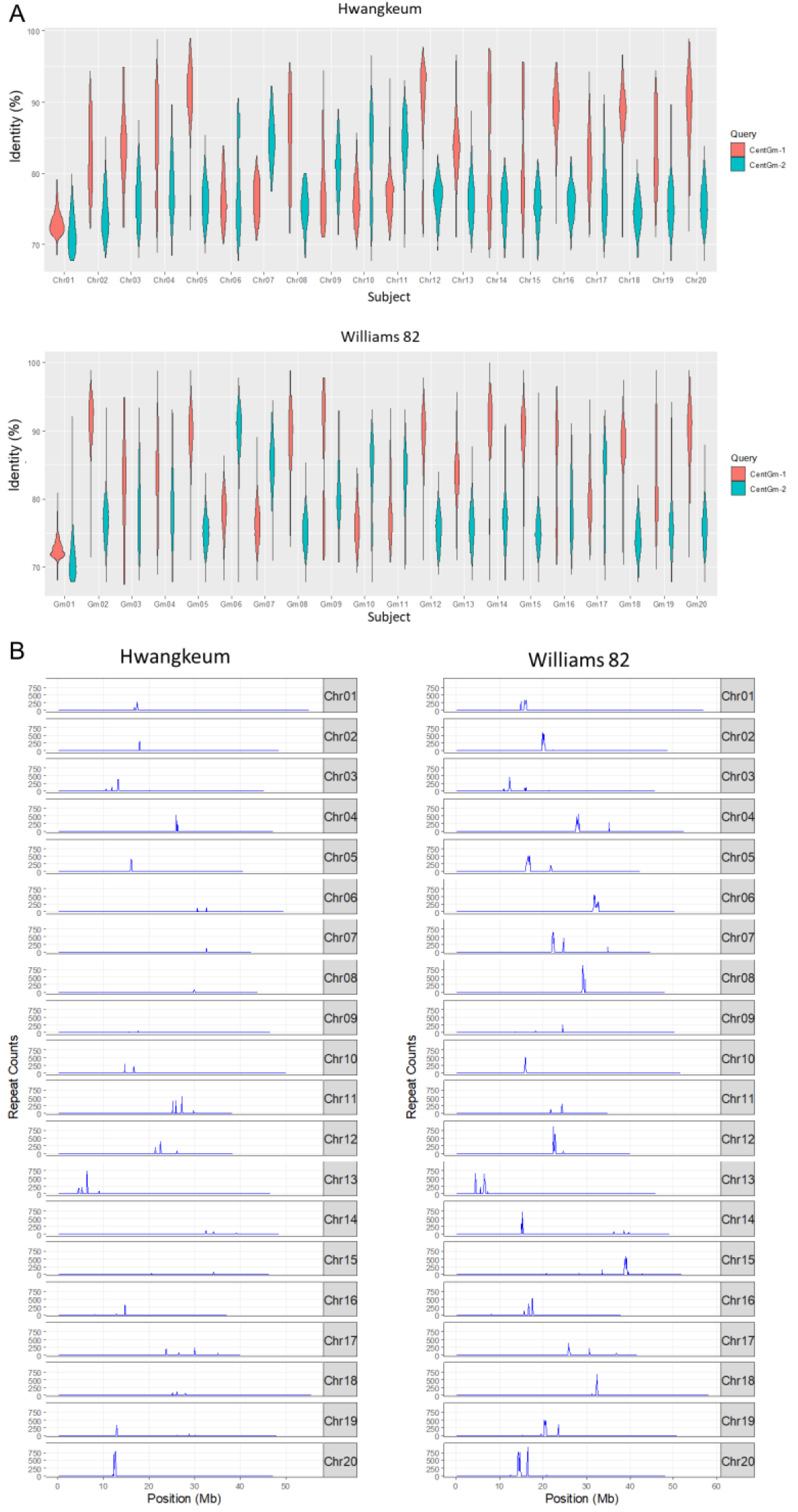
Genome-wide distribution patterns of centromeric repeats in the Williams 82 and Hwangkeum assemblies. (A) Violin plot distributions of the percent identity of centromeric repeats hit by BLAST searches with CentGm-1 and CentGm-2, respectively, along the 20 soybean chromosomes, as sampled in the Hwangkeum and Williams 82 assemblies. (B) Genome-wide centromeric repeat density in the Hwangkeum and Williams 82 assemblies. Centromeric repeats hit by CentGm-1 or CentGm-2 were combined by removing one of overlapping repeat sequences and then the repeat sequence density was plotted in 100-kb windows along the 20 soybean chromosomes.

Searches of structural variants (SVs) in the Hwangkeum assembly relative to the Williams 82 reference sequence resulted in 11,542 deletions (≥ 50 bp), 10,845 insertions (≥ 50 bp), 2,504 interchromosomal translocations (> 1,000 bp), and 168 inversions (> 10 kbp) (File S3). The total length of insertions (27.5 Mb) was 5.6 Mb longer than that of deletions (21.9 Mb). Our close examination suggested that the length difference was largely due to the insertions of unanchored scaffolds in the Williams 82 assembly. For example, most of scaffold_21 (3.57 Mb) and half of scaffold_22 (1.24 Mb), which are the two longest unanchored scaffolds in the Williams 82 assembly, were inserted with inverted orientation into chromosome 11. Scaffold_21 corresponded with the largest insertion of 3.46 Mb, and scaffold_22 corresponded with a cluster of several large (> 7 kb) insertions that were likely separated by repetitive sequences. Therefore, the insertion of scaffold_21 and scaffold_22, which was also predicted by our previous genetic mapping study (Lee *et al*. 2020), is the main cause of the size increase of chromosome 11 in Hwangkeum relative to the Williams 82 reference sequence. However, note that this is not a natural event and also indicates an improvement in the Hwangkeum assembly.

The sizes of the detected interchromosomal translocations ranged from 1,001 bp to 184,994 bp with median of 3,294 bp. When we searched for 109 putative misjoint chromosomal regions in the soybean Williams 82 reference genome sequence (Wm82.a2.v1), which required re-positioning to different chromosomes based on genetic maps constructed in our previous study (Lee *et al*. 2020), more than 80 regions were located at different chromosomes in the Hwangkeum genome, as predicted. The results demonstrate the soundness of our misjoint detection method, as well as the improvement in the Hwangkeum assembly. Those misjoint regions that required re-positioning by multiple markers tended to contain multiple adjacent blocks, and thus the adjacent blocks could be merged together to treat them as the same large misjoint event, in accordance with a previous method for human genome study (Audano *et al*. 2019). As expected, each of the merged blocks tended to correspond with a large indel longer than 100 kbp, thereby indicating evidence of another improvement in the Hwangkeum assembly. One exception is the movement of a 2.43 Mb fragment between the 36.99 Mb and 39.42 Mb positions from chromosome 15 in Williams 82 to chromosomes 4 (approximately 0.38 Mb), 5 (0.57 Mb), and 13 (1.48 Mb) in the Hwangkeum (Table S3). Although no markers were located on these chromosomal regions in the WH and HI maps, the fragment in the Williams 82 assembly is likely a concatenated scaffold. Interestingly, these putative artifacts explained the relatively larger decrease of chromosome size in chromosome 15 and slight increase in chromosome 13. Despite the gain of the ∼ 0.38 Mb fragment, an approximately 0.83 Mb fragment was translocated from chromosome 4 (Williams 82) to chromosome 3 (Hwangkeum), as predicted by the genetic mapping, thereby partly explaining the decrease in the length of chromosome 4. Taken together, our results suggest that the difference of the total lengths of insertions and deletions is not the main cause for the shorter total assembly length of the Hwangkeum assembly than that of the Williams 82 assembly.

The detected 168 inversions comprised 64 inversions and 104 intrachromosomal translocation & inversions (File S3). Among the predicted inversions, each of the 94 inversion fragments clearly matched with a single contig. Closer inspection of these inversion fragments indicated that because most of these contigs contained a single marker or multiple cosegregating markers in our WH and HI genetic maps, they could not be oriented in the ALLMAPS assembly process. Approximately 40 inversions that were part of a contig or covered by part of two contigs were located at low-recombination chromosomal regions, and so neither could their orientations be determined by genetic markers. At least 13 inversions were apparent errors by the ALLMAPS assembly because their orientations were inversed against the orders of the markers with one or two recombination events in the two genetic maps. All these putative artificial inversions were marked in the list of detected inversions (File S3). Which of the Hwangkeum or Williams 82 assemblies, both of which used genetic maps for pseudomolecule construction, contains correct orientations for these putative artificial inversions is unknown at this point because most of them locate at low-recombination chromosomal regions. Excluding all these putative assembly errors, 27 predicted inversions remained to be real. In the results, most of the detected inversions were not supported by genetic markers, and only 27 detected inversions appeared to be imbedded within a contig and the total length of the inversions was 1.86 Mb. Among the 27, seven were supported by genetic marker orders. The sizes of the 27 inversions ranged from 10 kb to 211 kb. As the genome-wide average recombination rate in soybean was estimated to be 2.5 cM/Mb (Lee *et al*. 2013), this result suggests that the inversions may not have a substantial impact on the genetic difference between Hwangkeum and Williams 82. The two largest detected inversions were adjacent but not overlapping 211-kb and 201-kb fragments between 31.89 Mb and 32.47 Mb positions on chromosome 7 in the Hwangkeum assembly. Although many of the breakpoint junctions of the detected inversions appeared to be located on repetitive sequences, we attempted to validate the two largest inversions by PCR-amplification using primers spanning their breakpoint junctions (Figure S2). Sequence comparison between the Hwangkeum and Williams 82 assemblies suggested that there might be some possibility generating specific primers from one side of the 211-kb inversion and from both sides of the 201-kb inversion. However, only one primer set, which was designed for amplification of one breakpoint junction of the 201-kb inversion, gave a specific PCR product that was subsequently confirmed by sequencing, supporting the correct assembly of the Hwangkeum genome.

### Diversity and evolution of centromeric satellite repeats

The differences of the locations, numbers, and ratios of the two repeats that distributed across soybean chromosomes supported the notion that differential distributions of these distinct repeats may reflect the allopolyploid nature of soybean (Gill *et al*. 2009), and then were used for the karyotyping of 20 soybean chromosome pairs (Findley *et al*. 2010). As we identified nearly nine times more satellite repeats from the Williams 82 assembly, we decided to further investigate the distribution patterns and evolution of centromeric repeats across chromosomes to investigate the integrity of the Hwangkeum genome assembly. We first compared the two groups of satellite repeats hit by BLAST searches with CentGm-1 and CentGm-2, respectively, from the Hwangkeum and Williams 82 assemblies (Figure 2A and Figure S3). The distribution patterns of percent identity values from the BLAST searches within each of the chromosomes could be divided into three groups: First, both CentGm-1- and CentGm-2-hit repeats showed lower than 80% identity (chromosomes 1, 4, 6, 9 and 19); second, the CentGm-1-hit repeats showed higher percent identity than the CentGm-2-hit repeats (chromosomes 2, 3, 5, 8, 12, 13, 14, 15, 16, 17, 18, and 20); and the CentGm-1-hit repeats showed lower percent identity than the CentGm-2-hit repeats (chromosomes 7, 10, and 11). The distribution patterns could also be divided into two groups of narrow or wide identity value distributions. Despite the large difference of the numbers of repeats identified, the distribution patterns were quite similar between the Williams 82 and Hwangkeum assemblies. The results suggested that the higher diversity of repeat sequences might not be due to assembly errors but reflect polymorphisms of repeats generated during the evolution of each chromosome. Interestingly, approximately half of the unanchored contigs that are assumed to be subject to much less degree of assembly errors showed wide identity value distributions (Figure S3).

The genomic distribution of the unique satellite repeats in 100-kb windows along the 20 soybean chromosomes showed that the centromere on each chromosome revealed different patterns of repeat density peaks (Figure 2B). Although the highest peaks of centromeric repeats between the two assemblies on most of the pseudomolecules corresponded to each other, the Williams 82 assembly showed more additional peaks. Notably, while the Williams 82 assembly showed two centromeric locations separated by more than 10 Mb from each other on chromosomes 7 and 14, Hwangkeum showed single locations on both the chromosomes. Five chromosomes (3, 4, 15, 19, and 20) in the Williams 82 assembly showed two centromeric locations separated by several Mb from each other. Separations of putative centromeric regions by more than 10 Mb were also observed on four chromosomes in the updated Zhonghuang 13 assembly (Shen *et al*. 2019). With some exceptions such as the point centromeres or holocentromeres, monocentric centromeres from plant to animal species are normally established on highly repetitive DNA arrays that usually contain distinct centromeric repeats (Cuacos *et al*. 2015; Barra and Fachinetti 2018). A fluorescent *in situ* hybridization study revealed the presence of monocentric centromeres across the soybean genome (Findley *et al*. 2010). Thus, the observation of more monocentric centromeres in the Hwangkeum assembly is evidence that despite the shorter total length of centromeres, the Hwangkeum assembly has been improved relative to the Williams 82 reference assembly in terms of overall scaffold order and position in the pericentromeric regions of the assembly.

### Phylogenetic analysis of centromeric satellite repeats

For phylogenetic analysis, repeat sequences < 89 bp or > 96 bp were removed from the combined set of 25,030 repeat sequences from the Hwangkeum assembly for the sake of alignment. The resultant 20,386 repeat sequences were aligned, and a Neighbor-joining distance tree was constructed. Four major clusters were found (Figure 3), in contrast to the previous report that there were two major groups of centromeric repeats in the soybean genome (Gill *et al*. 2009; Valliyodan *et al*. 2019). Because the representative repeat sequences previously reported belong to the two most distant groups, CentGm-1 group was renamed as CentGm-1a, and CentGm-2 as CentGm-2a. Of the two novel groups between CentGm-1a and CentGm-2a, the group next to CentGm-1a was referred to as CentGm-1b, and the group next to CentGm-2a as CentGm-2b. The finding of the two novel groups in this study was likely due to the fact that we used less stringent BLAST cut-off criteria with blast-short and gap penalty options, in addition to the cutoff of 60% sequence identity and 80% match length used in the previous studies. Interestingly, the observation of four repeat groups are somewhat consistent with the hypothesis that the differential distributions of soybean satellite repeats may reflect the allopolyploid nature of soybean (Gill *et al*. 2009).

**Figure 3.**
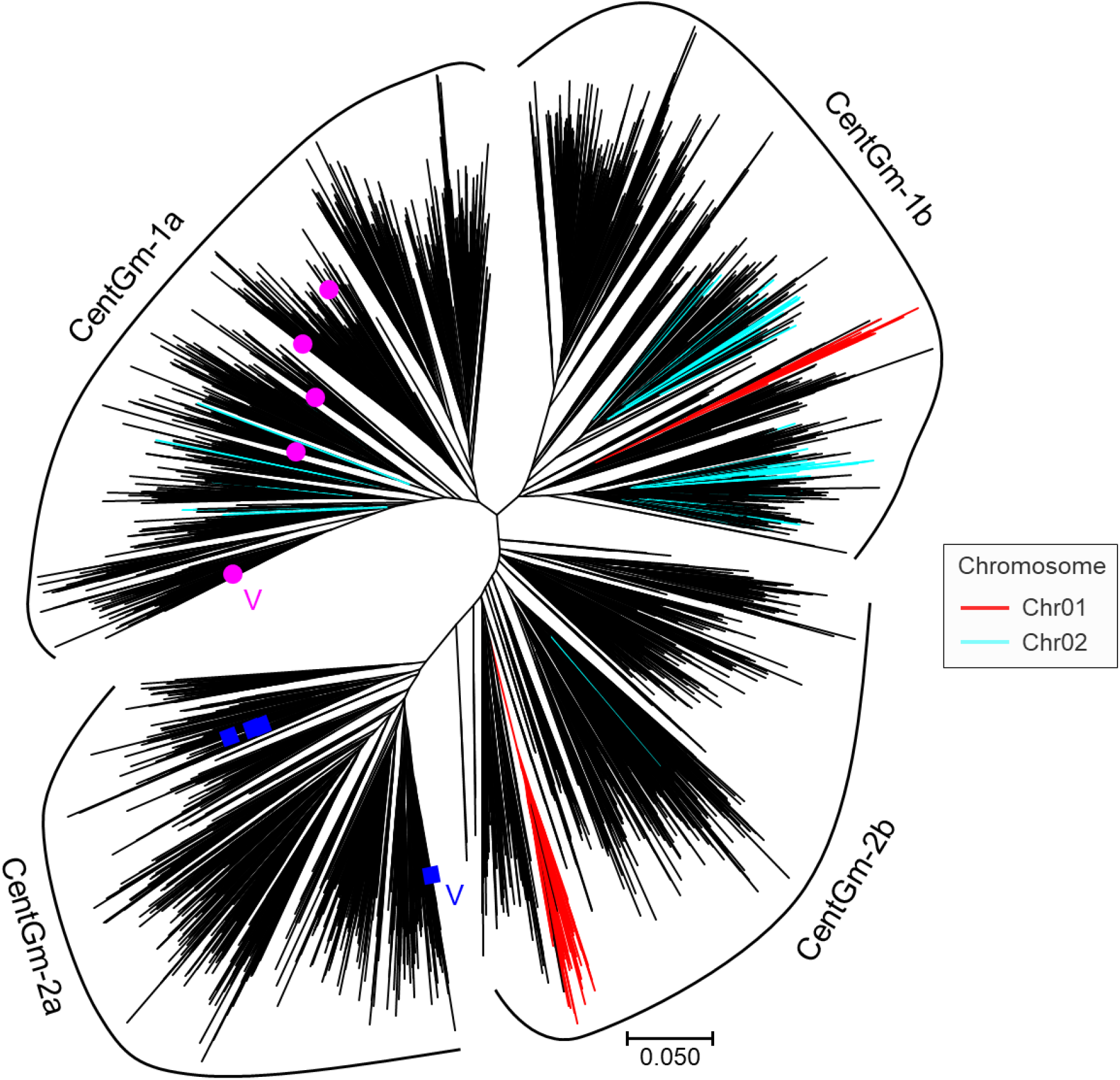
Neighbor-joining phylogenetic tree of 4469 centromeric repeat sequences in the Hwangkeum assembly together with nine publicly available representative sequences. Repeat sequences hit by BLAST searches with CentGm-2 or CentGm-1 were combined and then clustered with a cutoff of 90% similarity. Repeat clusters with lengths ranging from 88 to 95 bp were used for further analysis. Representative repeat sequences publicly available are indicated by pink circles for CentGm-1 and by blue squares for CentGm-1. The sequences used for BLAST searches were also highlighted by V. Centromeric repeat sequences were grouped into four subgroups; CentGm-1a, CentGm-1b, CentGm-2a, and CentGm-1a. Sequences on chromosome 1 are indicated by red branches and those on chromosome 2 by light blue branches.

Major portions of repeat sequences in each of the chromosomes appeared to belong to two adjacent groups, with exceptions of chromosomes 4 and 17 where the repeat sequences were spread over four groups and three groups, respectively (Figure 3 and Figure S4). Most of the chromosomes do not contain one or two of these four centromeric repeat groups. Dispersion of each of the four repeat groups on a number of chromosomes may represent relics of ancestral arrays rather than the mixing of chromosomes or assembly errors. This result indicates that rapid and dynamic changes in the centromeric DNA after the formation of the tetraploids may have occurred preferentially within each of the chromosomes rather than the intermixing of chromosomes. Thus, our result is somewhat consistent with significant genetic variation within centromeric satellites and asymmetrical distribution of centromere organization among the three subgenomes observed in hexaploid wheat (Lee *et al*. 2005), providing additional evidence for the integrity of the Hwangkeum assembly.

### Identification of centromeric satellite repeats in *Glycine latifolia*

The weakness or absence of hybridization with satellite repeats to genomic DNA within a genus suggested the rapid divergence of centromeric satellite repeats (Lee *et al*. 2005; Gill *et al*. 2009; Ta *et al*. 2021), and in the case of rice relatives, novel divergent satellite repeats with low or no sequence similarity with CentO were isolated from several relatives. As genome sequence of *G. latifolia* (Liu *et al*. 2018), a perennial relative of soybean, is available, we searched CentGm repeats in the *G. latifolia* genome. Interestingly, we extracted 3,107 non-redundant repeat sequences using CentGm-1 and CentGm-2. The percent identity of those sequences with CentGm-1 and CentGm-2 ranged from 67% to 83%, consistent with the previous Southern hybridization results (Gill *et al*. 2009). Examination of sequence regions containing *G. latifolia* repeat using the Tandem Repeat Finder indicated that most of the repeats are 91-bp monomer unlike the 91- or 92-bp monomers in soybean (File S4). Of the repeats detected by the Tandem Repeat Finder, 73 of the 90-bp repeats and 2,944 of the 91-bp repeats were members of the set of 3,107 repeats identified by the BLAST searches, and 92-bp repeats were absent in the 3,107 set.

The 3,107 repeat sequences were combined with five CentGm-1 representative sequences, four CentGm-2 representative sequences, and ten sequences from each of the CentGm-1b and CentGm-2b groups. The resultant 3,046 repeat sequences were aligned, and a Neighbor-joining distance tree was constructed (Figure S5). Interestingly, the diverse types of soybean sequences were clustered into one large group interspersed with *G. latifolia* repeat sequences. Unlike the sequence divergence between the 91-bp and 92-bp repeat units in soybean, the 90-bp repeat sequences were also interspersed with 91-bp repeat sequences. The results indicated that although further investigation will be required because the *G. latifolia* assembly contained a much lesser number of repeats than the Hwangkeum or Williams 82 assemblies, *G. latifolia* genome likely contains significantly divergent CentGm-type centromeric satellite repeats, reflecting the evolutionary distance between the two species. Nevertheless, observation of a unique repeat group in the *G. latifolia* assembly might provide an opportunity to further test the hypothesis that differential distributions of soybean satellite repeats may reflect the allopolyploid nature of soybean (Gill *et al*. 2009).

### Annotation of the Hwangkeum genome and gene content comparison with other publicly available soybean genomes

Repetitive sequences made up 50.2% of the Hwangkeum genome (Table 1 and Table S7). Long terminal repeat (LTR) transposable elements were the most abundant elements (83.8% of repetitive content), including the Gypsy (56.7% of repetitive content) and Copia (26.3% of repetitive content) families. The portion of the repetitive sequences in the Hwangkeum genome appeared to be lower than the 60.6% of the Williams 82 genome, which was likely overestimated, and the average of 54.5% of the 26 soybean genomes assembled using PacBio sequencing data. Even if the satellite tandem repeats (∼ 1.0%) detected in the 26 soybean genomes are excluded, the Hwangkeum genome contained at least 3% (approximately 30 Mb) lower amount of repetitive sequences than the reported soybean genomes. In addition to the collapse of centromeric satellite repeats described above, this result suggests that the assembly collapse of repetitive sequences is likely a main cause of the shorter total lengths of the Hwangkeum assemblies relative to those of the Williams 82 and other soybean assemblies (Tørresen *et al*. 2019).

A total of 79,870 transcripts for 58,550 protein-coding genes were found, which numbers are comparable to 88,647 transcripts for 61,303 genes in the reference soybean genome Wm82.a2.v1 (86,256 transcripts for 52,872 genes in the updated Wm82v4 assembly). Assessment of the annotation completeness with two BUSCO databases, eukaryota odb10 and embryophyta odb10, indicated that the gene content was effectively captured in the PromethION assembly (Table S8): BUSCO analysis against eukaryota odb10 and embryophyta odb10 demonstrated 247/255 (96.9%) and 1,562/1,614 (96.8%) of BUSCO genes from the assembly, respectively. Of the 79,870 transcripts, 76,823 (96.2%) were associated with EggNOG functional categories (Table S9), 56,212 (70.4%) had an InterPro match, 56,682 (71.0%) had a PFAM match, and 40,345 (50.5%) were assigned a gene ontology (GO) term (File S5). We annotated 327 NLR genes, the genes of agronomically important superfamily, in the Hwangkeum assembly using the Seqping pipeline, which number is much lower than the 477 in the Williams 82 Wm82.a2.v1 assembly. As TGFam-Finder was recently used to annotate 66 additional NLR genes from the Williams 82 Wm82.a2.v1 assembly (Kim *et al*. 2020), we re-annotated the NLR genes using TGFam-Finder in the Hwangkeum assembly. A total of 503 NLR genes were annotated using TGFam-Finder in the Hwangkeum assembly with 176 additionally predicted genes (File S6), resulting in a similar number of annotated NLR genes between the Hwangkeum and Williams 82 assemblies.

A total of 26,433 orthologous groups were identified between the Hwangkeum and Williams 82 assemblies using OrthoMCL. The Hwangkeum and Williams 82 assemblies possessed 24,977 and 25,445 orthologous groups, respectively. Of them, 23,989 orthologous groups (90.7%) existed in common between the Hwangkeum and Williams 82 assemblies. With the same criteria, about 4.0% of the Hwangkeum genes (988) and about 5.7% of the Williams 82 genes (1,456) were lineage-specific orthologous groups in the Hwangkeum and Williams 82 genome, respectively. The portions of lineage-specific genes, which are dispensable genes in terms of pan-genome, are somewhat lower than those of the recent soybean pan-genome analysis (Liu *et al*. 2020) that showed that dispensable gene families accounted for an average of 19.1% of the genes in individual accessions. Thus, this result indicates a close relationship between Hwangkeum and Williams 82.

Finally, to test the quality of the Hwangkeum assembly down to the nucleotide level in the euchromatic regions, we examined the presence of known polymorphisms at genetic loci associated with golden seed color and strong SMV resistance, which are two characteristics of Hwangkeum, and whose genes have recently been characterized (Chen *et al*. 2002; Yang *et al*. 2010; Jeong and Jeong 2014; Redekar *et al*. 2016). To characterize seed coat and flower colors, Yang *et al*. (2010) developed 28 markers from eight enzyme-encoding gene families and a transcription factor that had been characterized as regulating anthocyanin biosynthesis or were homologous to the genes characterized in other plants. Those markers were mapped in a Hwangkeum by IT182932 population. We confirmed that Hwangkeum polymorphic sequences of the 28 markers were present in the Hwangkeum assembly at the chromosomal locations predicted by both the genetic mapping as well as the Williams 82 assembly (Table S10). Thus, the results provide evidence for the high quality of the Hwangkeum assembly.

Hwangkeum is resistant to SMV, while Williams 82 is susceptible to SMV. The high level of resistance to all SMV strains in Hwangkeum was initially ascribed to a single dominant *Rsv*1 allele (Chen et al., 2002). However, Jeong and Jeong (2014) found that Hwangkeum contains more than two resistance genes at the classical *Rsv*1 locus as well as the *Rsv*3 locus. The two loci act in a complementary manner, in which the *Rsv*3 locus tends to confer resistance to SMV strains that are virulent to *Rsv*1-carrying plants. This locus is also interesting because it is located in the middle of a heterogeneous cluster (Suh *et al*. 2011) that contain members of the NLR as well as leucine-rich repeat receptor-like kinase (LRR-RLK) multigene families, of which some members have been reported to be disease resistance genes (Song *et al*. 1997; Parniske and Jones 1999). A strong candidate *Rsv*3 gene was proposed by a comparative sequence analysis (Redekar *et al*. 2016), and was then validated by overexpression and transient silencing (Tran *et al*. 2018; Ross *et al*. 2021). When the gene arrangement at this complex region spanning 1.83 Mb delimited by sequence-based markers Satt063 and GSINDEL133985 (Lee *et al*. 2013) was compared between the Hwangkeum and Williams 82 assemblies, the order and orientation of the shared genes were remarkably consistent with each other. Twenty-four of the 184 genes were unique to the Williams 82, and 12 of the 24 unique genes appeared to be functionally unannotated. In the case of the Hwangkeum assembly, 25 of the 168 genes were unique, and 23 of the 25 appeared to be functionally unannotated. Thus, those unique genes might have resulted from over-annotation of either assembly. The smaller total number of genes in the Hwangkeum is likely due to poor annotation in the multigene tandem repeat cluster by the Seqping pipeline because the TGFam-Finder added three more NLR genes at the *Rsv*3 locus. When the arrangement of only the NLR and LRR-RLK genes were examined between the two assemblies at this *Rsv*3 region, the order and orientation of the genes were consistent with each other, as we highlighted homologs of the cloned *Rsv*3 gene (Figure 4). The Williams 82 assembly contained one more partial NLR gene and one more LRR-RLK gene relative to the Hwangkeum. Interestingly, the Williams 82 Wm82.a2 version contained five LRR-RLK genes, while the Williams 82 Wm82.a1 version contained 10 LRR-RLK genes in our previous study (Suh *et al*. 2011), thereby indicating the much improved assembly in the Wm82.a2 version. Therefore, the high similarity of gene arrangement between the Hwangkeum and Williams 82 assemblies suggests that the gene-rich euchromatic regions of the Hwangkeum assembly are of a similar quality to those of the Williams 82 soybean reference genome sequence at the nucleotide level

**Figure 4.**
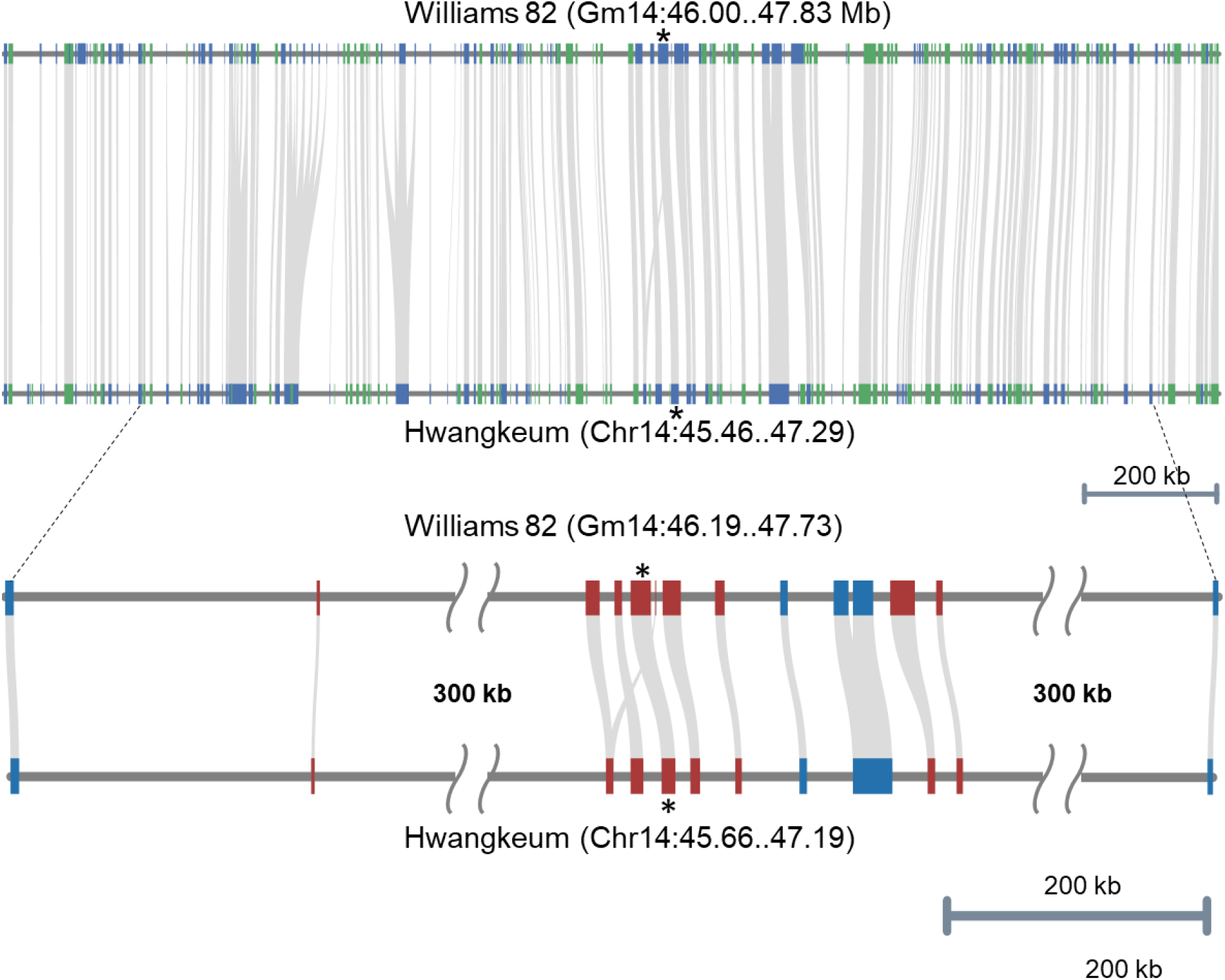
Comparison of gene arrangement between the Hwangkeum and Williams 82 assemblies at the chromosome 14 region in the vicinity of the *Rsv*3 locus. A. Comparison of order and orientation of all homologous genes between the Williams 82 and Hwangkeum assemblies. Genes are indicated by blue and green boxes in an alternate manner. Homologs of the cloned *Rsv*3 genes are indicated by asterisk. B. Comparison of order and orientation of nucleotide-binding and leucine-rich-repeat (NLR) genes and leucine-rich repeat receptor-like kinase (LRR-RLK) genes that show a heterogeneous cluster. NLRs are indicated by red boxes and LRR-RLKs by blue boxes.

## Conclusions

In this study, we report the *de novo* assembly of the palaeopolyploid soybean genome through the integration of genetic linkage mapping and Nanopore PromethION sequencing. The total length of the present assembly (931 Mb) was shorter than that of the PacBio SMRT assembly of Hwangkeum (966 Mb) in this study as well as those of the public data (> 970 Mb). The shorter assembly length is likely caused by assembly collapse at repeat regions (Tørresen *et al*. 2019), including centromeric satellite repeat regions as well as transposon repetitive sequences. However, several lines of evidence have suggested that the assembly quality of Hwangkeum at the chromosome level was more improved than the public assemblies. First, our enhanced detection of centromeric satellite repeats that resulted in a much greater number of repeats and the finding of two novel repeat groups revealed more monocentric centromeres across all 20 chromosomes, which is consistent with the chromosomal nature of soybean genome predicted by the fluorescent *in situ* hybridization study (Findley *et al*. 2010), in the Hwangkeum assembly relative to the Williams 82 assembly. Second, we demonstrated that much shorter chromosomes or longer chromosomes could be explained by the predicted misjoints or insertions of unanchored scaffolds in the Williams 82 assembly, most of which were predicted by our previous genetic map study (Lee *et al*. 2020). Moreover, genetic markers or cloned genes associated with golden seed color and strong SMV resistance were located as predicted by previous genetic studies in the assembled chromosomes of Hwangkeum and the order and orientation of the examined genes were remarkably similar between the Hwangkeum and Williams 82 assemblies. Importantly, the BUSCO analyses indicated that the genome sequence and gene content qualities of our Hwangkeum assembly are comparable to those of the public assemblies. Thus, both the examinations of gene contents at genome-wide and specific chromosomal regions as an evolutionary measure of genome completeness suggest that the Hwangkeum assembly is a high-quality assembly. Different sequencing technologies show different pros and cons in the genome assembly projects (De Maio *et al*. 2019). Consequently, the present study shows that *de novo* genome assembly using the Nanopore PromethION long-reads platform provides promising results. Thus, this high-quality genome assembly for Hwangkeum will facilitate genetic dissection of the distinctive organoleptic and agronomical features of Hwangkeum, one of the typical cultivars in the Korean climate, as well as a better shaping of the soybean pan-genome.

## Acknowledgments

This work was supported by the National Research Foundation grant (NRF-2018R1A2A2A05021904) funded by the Korea government awarded to SCJ and by the Korea Research Institute of Bioscience and Biotechnology Research Initiative Program

